# A role for the *Fem-1* gene of *Drosophila melanogaster* in adult courtship

**DOI:** 10.1101/2020.01.19.911693

**Authors:** Miles Thies, Brett Berke

## Abstract

The *Fem* family of genes influences sex determination and/or the development of sex-specific characteristics in a wide variety of organisms. Here, we describe the first mutational analysis of the *Fem-1* gene of *Drosophila melanogaster*. The amino acid sequence of the two *Drosophila Fem-1* transcripts are moderately conserved compared to that of both *Fem-1* in *C. elegans* and the two *Fem-1* transcripts in humans, with multiple ankyrin repeats. Using two transposon-induced mutations of *Drosophila Fem-1*, we observed striking defects in adult courtship behavior that are attributed to defects in male courting as opposed to female receptivity. Specifically, viable *Fem-1* mutant males courted *Fem-1* females more vigorously with an increased amount of chasing and singing than pairs of control flies. Nevertheless, *Fem-1* males did not copulate at a higher frequency than controls. The above courtship defects persisted when *Fem-1* males courted control females, but no phenotypes were observed when control males courted *Fem-1* females. These results indicate that *Drosophila Fem-1* may interact with other genes involved in courtship and sex determination. *Fem-1* mutants also suppressed wing and body growth, consistent with the actions of a homologue in mice. Additional analyses of these *Fem-1* alleles will help address the nature of these mutations, deepen our molecular understanding of courtship, and contribute to the evolutionary relationships among this highly conserved gene family.

## Introduction

The *Fem-1* gene was first identified in *Caenorhabditis elegans*, where it plays a vital role in the development of male worms (Doniach & Hodgkin, 1984) through the ubiquitination and degradation of sex-determining proteins (Chan et al., 2000; Spence et al., 1990; Starostina et al., 2007). The *Fem-1* gene family is highly conserved across a variety of animal phyla, and it is implicated in sexual development in porifera (Perović-Ottstadt et al., 2004), arthropods (Galindo-Torres et al., 2019; Koch et al., 2014; Ma et al., 2012; Montana & Littleton, 2006; Rahman et al., 2016; Shulman & Feany, 2003), mollusks (Teaniniuraitemoana et al., 2014), and chordates (Chan et al., 2000; Galindo-Torres et al., 2019; Gilder et al., 2013; Krakow et al., 2001; Lu et al., 2005; Oyhenart et al., 2005; Qin et al., 2019; Shi et al., 2011; Ventura-Holman & Maher, 2000; Ventura-Holman et al., 1998; Ventura-Holman et al., 2003; Wang et al., 2008). Fem-1 proteins are highly expressed in neural tissues (Ventura-Holman & Maher, 2000) and have also been implicated in a few neuronal processes. *Fem-1B* mutations in mice disrupt the insulin signaling that is critical for neuronal growth (Lu et al., 2005). The mouse hippocampus increases expression of Fem-1C in response to ischemia (Jin et al., 2001). Fem-1 also modulates neurodegeneration caused by over-expression of the Tao protein in a *Drosophila* model of Alzheimer’s disease (Shulman & Feany, 2003). In non-neuronal contexts, Fem-1b modulates the Gli1 transcription factor and interacts with Ankrd37 in cell culture (Gilder et al., 2013; Shi et al., 2011), and binds to the Nkx3.1 transcripition factor and the Phtf1 ER protein in mouse testes (Oyhenart et al., 2005; Wang et al., 2008).

Despite its strong evolutionary conservation and its expression in the fruit fly *Drosophila melanogaster*, the *Fem-1* gene of flies has never been examined for a role in adult courtship behaviors. The fly is a versatile model to understand the genetic basis of courtship because of its short life cycle, the ease with which flies can be maintained, and the extensive collection of mutations and genetic tools. Courtship behaviors produced by male flies are fixed action patterns (Villella & Hall, 2008), as they are genetically determined and relatively invariant between wild-type (WT) flies. When genetic mutations disrupt this stereotypy, the underlying causes can sometimes be traced back to effects on central neuron development (Yamamoto & Koganezawa, 2013). Genetic dissection of adult courtship indicates that a complex hierarchy of sex determination genes regulates sex-specific neuronal development and behavior (Yamamoto et al., 2014).

Courtship begins when the male fly orients his body towards the female. He may then tap her with a foreleg, sing to her by vibrating one wing (called a courtship song), chase after her, and lick her genitalia. Throughout this process, the female runs away from the male, but if she is eventually receptive, she will allow copulation. Mutational analyses have identified novel genes involved in courtship, helping to link alterations in neural circuitry with changes to the courtship program (Demir & Dickson, 2005; Finley et al., 1997; Shirangi et al., 2013, 2016; Zanini et al., 2012). Mutational analyses can also uncover the neurons that are necessary for distinct elements of the fixed action pattern of courtship behaviors (Kimura et al., 2008). While adult courtship has been extensively studied, recent findings indicate a surprising amount of complexity left to discover. For example, courtship behaviors may be influenced by circadian control (Fujii et al., 2017) and this fixed action pattern is sensitive to a variety of modulators (Ellendersen & von Philipsborn, 2017; Kim et al., 2017).

Here, we characterize the *Fem-1* gene in adult courtship behavior. We studied the effects of two *Fem-1* alleles and found that these mutants court more intensely that controls, without any change in copulation frequency. The mutants also disrupted the growth of the fly wing and body. Our phenotypic analysis of *Fem-1* therefore indicates an evolutionarily conserved role in sex determination and growth. The results lay a foundation for understanding how *Fem-1* interacts with well-studied courtship genes and the molecular mechanisms of the courtship phenotypes.

## Material and Methods

### Genetics

All fly stocks were raised at room temperature (about 21℃). The *0166-G4* (*w^1118^;PBac{IT.GAL4}Fem-1^0166-G4^*) and *EP 2065* (*w^1118^;P{EP}Fem-1^EP2065^*) stocks were obtained from the Bloomington Drosophila Stock Center (NIH P40OD018537; Department of Biology, Indiana University, Bloomington, IN, USA). The transposable element for the *EP 2065* allele is inserted into the 5’ UTR of the Fem-1a transcript and the insert for the *0166-G4* allele is located within the first intron of the Fem-1a transcript. The *w^1118^* stock was used as the genetic control for the two mutant alleles since both transposons were inserted into this genetic background. Amino acid sequence alignment of *Fem-1* proteins was done using the online multiple sequence alignment tool Clustal Omega (Sievers et al., 2011).

### Courtship assay

Single choice courting assays were performed at room temperature in a courting chamber made from plastic well plates (cut to 3mm in depth, 9mm diameter) covered with a glass coverslip. Courting chambers were lit from beneath using a lightbox. Before a courtship assay, the chamber was washed with 90% ethanol, left to dry for 5 minutes, washed with distilled water, and dried again. Male flies were collected 0-4 hours after eclosion and stored individually in vials with fly food for 4 days. Newly-eclosed female virgins were identified by the presence of a meconium and were stored at up to 10 virgins per vial with fly food for 4 days. For each courtship assay, a male fly was introduced into the courting chamber using a mouth aspirator and left to acclimate for 5 minutes. A female fly was then introduced into the chamber with the aspirator and the pair was then observed for 10 minutes. A camcorder (Sony HDR-CX405; Sony, New York, NY) was used to collect video recordings that were later analyzed by eye and using a MATLAB program (MathWorks, Natick, MA).

### Courtship analysis

Video recordings of courtship assays were manually reviewed and times were noted when the male fly was interacting, singing, chasing, or copulating. Interacting was a broad category used for any time the male was orienting, tapping, or licking, as it was generally difficult to distinguish between these individual behaviors. Courtship initiation was defined as the first instance in which the male engaged in any courtship behavior. The courtship vigor index was defined as the fraction of time the male spent interacting, chasing, or singing from initiation until successful copulation or the end of the 10 min observation period. The singing/chasing index was defined as the fraction of time the male spent singing/chasing during the entire observation period. The copulation percentage for each allele was defined as the percentage of mating pairs that initiated copulation during the observation period. Statistical comparisons of courtship indices were done using a Welch-t test assuming unequal variances in Microsoft Excel (Microsoft Corporation, Redmond, WA). The copulation percentages for each allele were compared using a chi-squared test in Microsoft Excel.

### Adult fly path length analysis

Video recordings of courting flies were analyzed using a MATLAB script (available upon request) that determined the total distance travelled by the courting flies during the observation period. This MATLAB script is based upon a previous video analysis system (Iyengar et al., 2012). During analysis, a graphical user interface prompts a user to input a region of interest, initial coordinates, and an intensity threshold for the conversion of each frame into a black-and-white image. The program then iterates through all frames, calculates the centroid of both flies, and identifies the male and female centroid by minimizing the distance travelled by each fly since the last frame. Once the analysis is complete, the user is then prompted to input the coordinates of the male fly for all frames where the sexes couldn’t be determined due to the flies overlapping in preceding frames.

### Fly Size Analysis

Before beginning these experiments, new fly stocks of all strains were made to ensure that all flies developed in similar environments. Briefly, adult flies were collected up to 6 hours after eclosion, separated based on sex, and then stored in vials with fly food for up to 4 days. For the *w^1118^* and *0166-G4* stocks, 15 males and 15 females were transferred to a new vial with fly food and a small, autoclaved piece of paper towel. Because the *EP 2065* stock was generally less healthy than the other stocks, 20 males and 20 females were used for this strain. About 10 days after these stocks were set, new adult flies began to eclose. These flies were collected up to 6 hours after eclosion, separated based on sex, and then stored in vials with fly food for 2-3 days. The flies were then anesthetized using diethyl ether (Fischer Scientific). Photos of the flies’ wings were taken after their removal. The flies were also arranged with their anterior side facing up and photos of the flies’ bodies were captured. These photos were analyzed using an open source MATLAB script to measure the length and area of the flies’ wings and bodies (www.mathworks.com/matlabcentral/fileexchange).

## Results

### Drosophila Fem-1 and its evolutionary conservation

The *Drosophila Fem-1* gene (Figure 1A) encodes two uncharacterized proteins: Fem-1a and Fem-1b. Both of these proteins (Figure 1B) have ankyrin repeat-containing domains, which mediate protein-protein interactions (Li et al. 2006). The percent identity between *Drosophila* Fem-1 and its homologous proteins in *C. elegans*, humans, and mice shows moderate conservation in amino acid sequence throughout the entirety of the protein (Figure 1C). The *Fem-1* alleles used in this study (*EP 2065* and *0166-G4*) result from transposons inserted near the N-terminus of the gene (Figure 1A), and their effects on the Fem-1 mRNA and protein are unknown.

**Figure 1.**
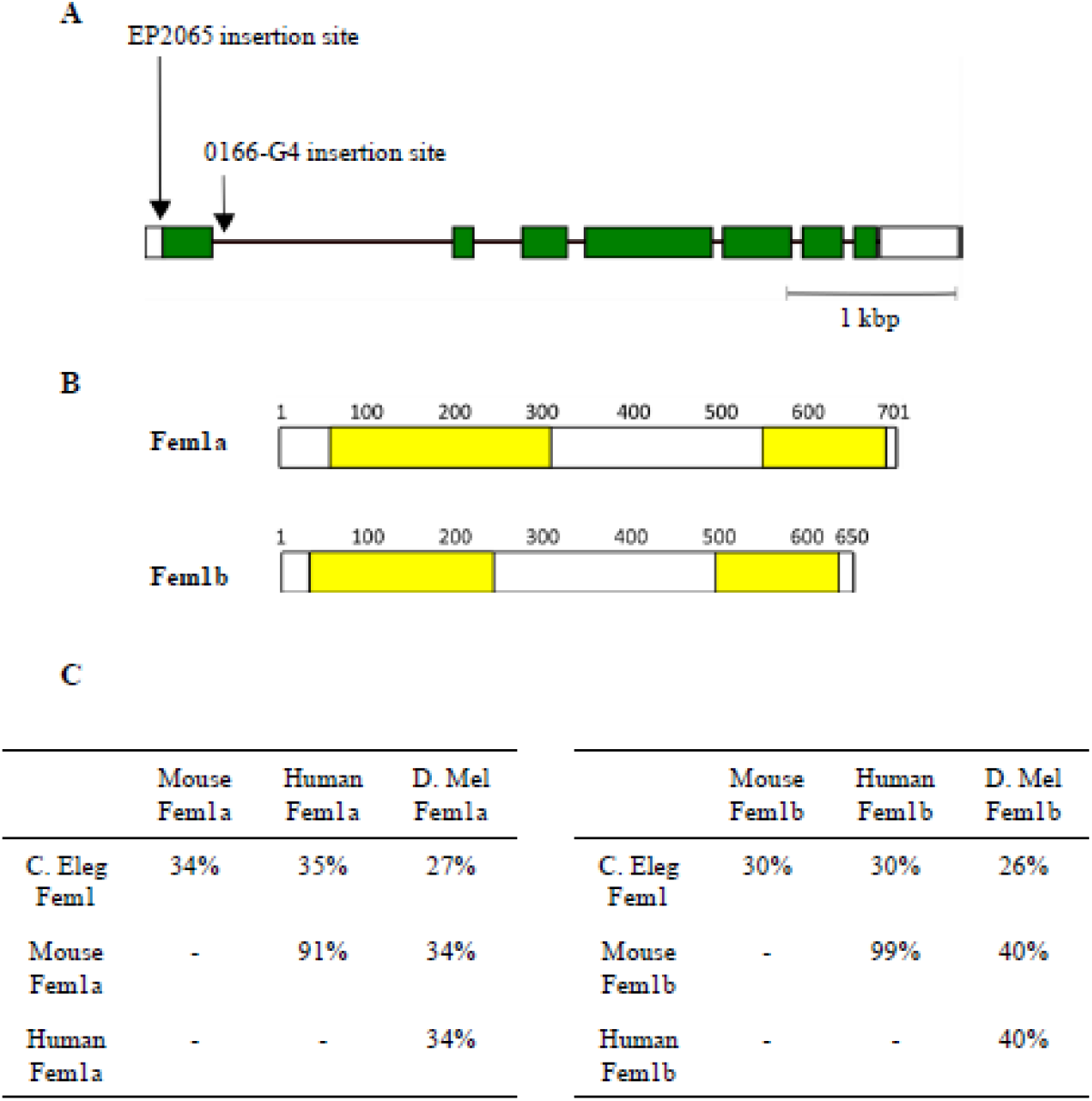
The *Fem-1* gene encodes a conserved protein in *Drosophila*, *C. elegans*, humans, and mice. (A) Model of the *Fem-1* gene in *Drosophila*. Exons are shown in green and untranslated regions in white. The insertion site of the transposable elements is shown for the two *Fem-1* alleles used here: *EP2065* and *0166-G4*. (B) Model of the Fem-1a and Fem-1b proteins in *Drosophila*. Ankyrin repeat-containing domains, which are known to mediate interactions with other proteins, are shown in yellow. (C) Percent identity matrix for Fem-1 proteins in *Drosophila, C. elegans*, humans, and mice.

### Mutations in Fem-1 result in increased courtship intensity with no change in copulation rate

Courtship assays were performed with *0166-G4* male/female, *EP 2065* male/female, and control *w^1118^* male/female pairs. These experiments used previously-isolated male and female virgin flies that were aged 4 days and introduced separately into a small mating chamber. Their stereotyped courtship behaviors over 10 min were then videotaped and analyzed by eye. Sample frames from these videos show orienting, chasing, singing, and copulating flies within the courting chamber (Figure 2). Replay of these videos was used to compute indices for singing and chasing, a courtship vigor index, and a latency to courtship. The measurements (see Methods) were used to characterize the amount of time flies exhibit singing and chasing, two behavioral elements of the courtship repertoire. The courtship vigor index indicates a broader set of behaviors (orienting, tapping, singing, chasing, licking), and together with the latency to courtship, gives a sense of the male’s drive to court (Krstic et al., 2009).

**Figure 2.**
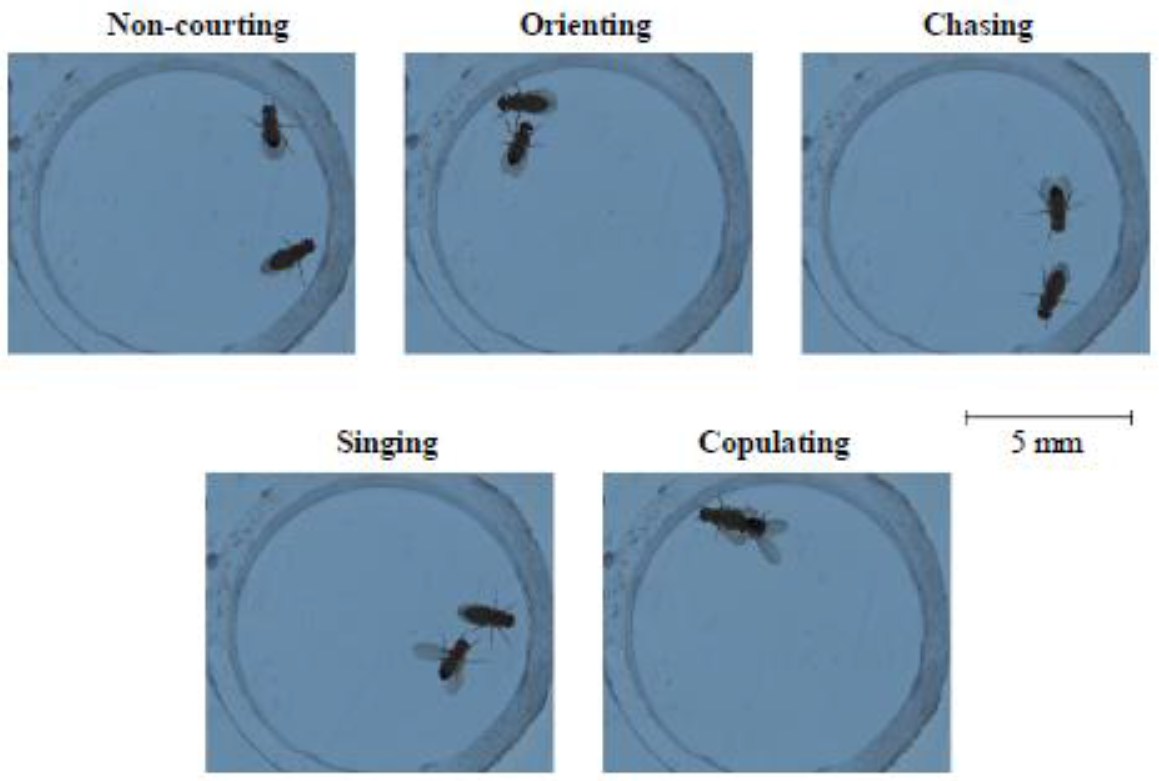
Representative images of adult courtship in *Drosophila*. Orienting: the male fly will approach and orient its body towards the female. Chasing: the male will chase behind the female. Singing: the male will stretch out and vibrate one wing while orienting towards the female. Copulating: the male will mount the female and complete copulation.

Comparisons between the three genotypes revealed a large increase in indices for singing (p = 4.8×10^−9^) and chasing (p = 3×10^−12^) for *0166-G4* pairs and a slight increase for *EP 2065* pairs (singing, p = 5.1×10^−3^; chasing, p = 8×10^−4^; Figure 3A-D). The *0166-G4* allele showed a significant increase in the mean courtship vigor index (p=9.4×10^−11^), but showed a non-significant trend of shorter mean latencies to courtship initiation. The *EP 2065* allele did not present any significant changes in mean courtship vigor index or latency to courtship. Given the increased courtship observed in both alleles, it was surprising that neither showed a significant increase in the percentage of mating pairs that copulated (Figure 3E). To address the possibility that changes in courtship resulted from changes in overall activity, a MATLAB program was created to measure the distance that each fly walked during the 10min courtship assay. Recordings where the mating pair successfully copulated were not used, as the flies stop moving once copulation begins. On average, the female fly moved a larger distance than the male fly for all genotypes. Both *Fem-1* alleles showed an increase in distance travelled for both sexes as compared to w^1118^ (EP 2065: male, p = 5×10^−6^; female, p = 8×10^−7^; *0166-G4*: male, p = 6×10^−16^; female, p = 5×10^−14^). It is difficult to determine if the increased courtship intensity in *Fem-1* mutants was caused by an overall increase in movement. While the changes in latency until initiation and chasing index could have been influenced by an overall increase in activity, the increased singing index in *Fem-1* mutants suggests that there were alterations to the neural circuitry for courtship.

**Figure 3.**
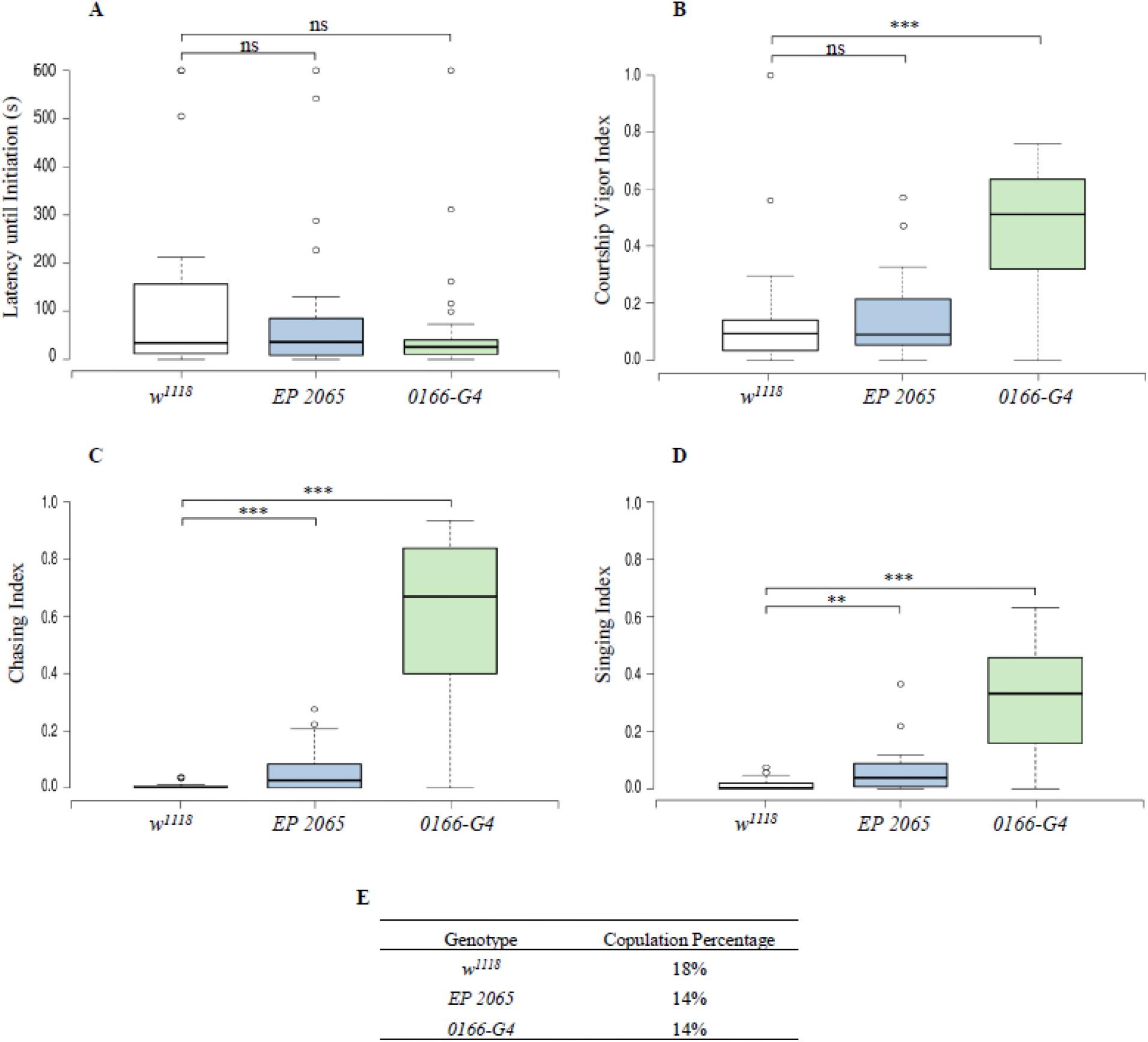
*Fem-1* mutants show an increase in courting intensity, including singing and chasing, without a similar increase in frequency of copulation. (A) Box plot depicting the mean latency to the initiation of courtship behaviors for the control and the two *Fem-1* alleles. Males that did not initiate courtship were assigned a latency of the entire observation period (600 s). There is no change in the mean latency to initiation between control and *Fem-1* mutants. (B) There is a significant increase in mean courting intensity in the *0166-G4* allele. (C) There is a significant increase in mean chasing index for both mutant alleles. (D) There is a significant increase in mean singing index for both mutant alleles. (E) There is no change in percentage of pairs that copulated between the three alleles. For details on how the intensities and indices were calculated, see the Methods. For *w^1118^*, n = 28; *EP 2065*, n = 28; *0166-G4*, n = 29 pairs.

Given that the loss of *Fem-1* alters the development of external sexual characteristics in *C. elegans (Doniach and Hodgkin, 1984)* and insulin signaling in mice *(Lu et al., 2005)*, we looked for gross structural abnormalities in *Fem-1* mutant male and female flies. Consistent with their ability to mate and their enhanced but otherwise normal courtship preference, we did not observe the loss of sex-specific structures or the switching of external genitalia (Figure 4). The apparently normal development of external sexual structures does not necessarily rule out a role for Fem-1 in these tissues, but it indicates that the courtship phenotypes in *Fem-1* mutants were likely caused by changes in development of the courtship circuitry within the CNS.

**Figure 4.**
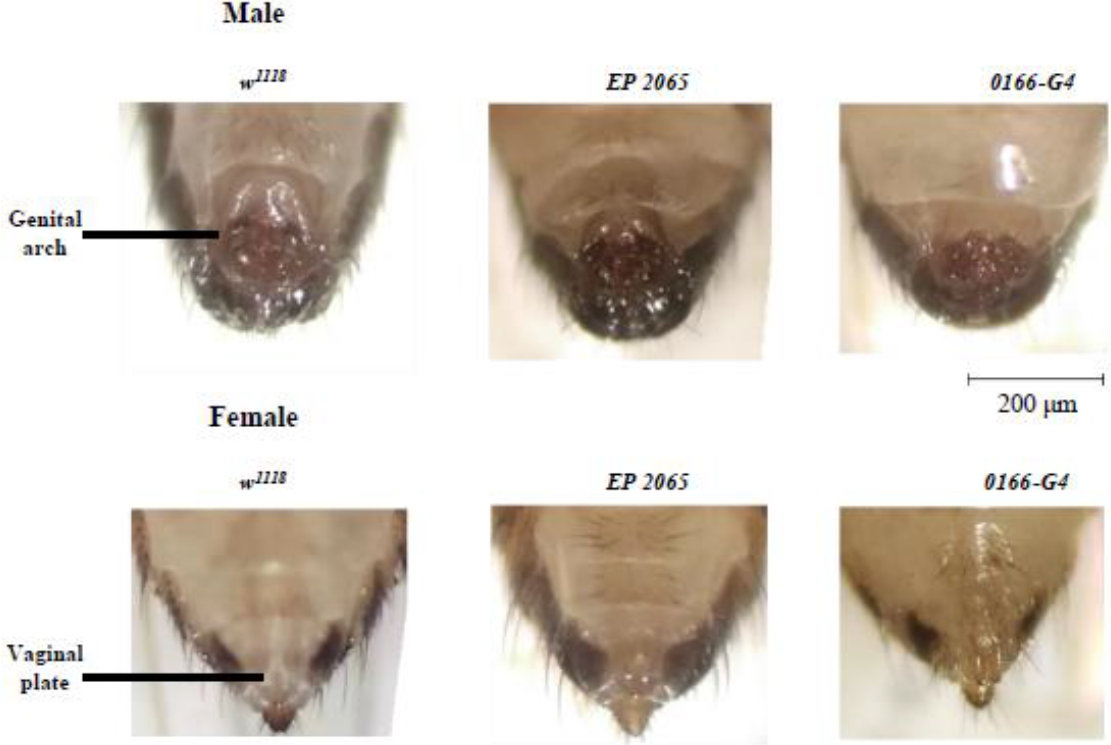
*Fem-1* mutations do not show obvious changes to the external genitalia of adult flies. The genital arch of the male and vaginal plate of the female are identifiable in *w^1118^* and *Fem-1* mutants. For *w^1118^*: males, n = 12, females, n = 16; *EP2065*: males, n=23, females, n=21; *0166-G4:* males, n=19, females, n = 20.

### Some Fem-1-dependent changes in courtship may be sex dependent

The characteristics of mating that were quantified in the above data were collected from mutant males, but it is possible that *Fem-1* mutations altered female receptivity, which in turn could affect male behavior. For example, if *Fem-1* females were less receptive while the males mated more vigorously, this might explain the unchanged copulation frequency. We therefore examined whether *Fem-1* mutations differentially affected male vs female flies. We performed courtship assays with *0166-G4* males / *w^1118^* females and *w^1118^* males / *0166-G4* females, as this allele showed the most striking courtship phenotype. Pairs of *0166-G4* male / *w^1118^* female flies had similar defects to *0166-G4* pairs (Figure 5). Their courtship characteristics differed significantly from *w^1118^* pairs and *w^1118^* male / *0166-G4* female pairs for many of the courtship parameters (compared to *w^1118^:* mean courtship vigor index, p = 3×10^−9^; mean chasing index, p = 8×10^−8^; mean singing index, p = 9×10^−7^; mean latency until initiation, p = 0.03; compared to *w^1118^* / *0166-G4*: mean courtship vigor index, p = 5×10^−10^; mean chasing index p = 8×10^−8^; mean singing index p = 6×10^−7^; and mean latency until initiation, p =0.06; Figure 5A-D). In contrast, the *w^1118^* male / *0166-G4* female pairs were only slightly different from *w^1118^* pairs (courtship vigor index, p = 0.018; singing index, p = 0.026; chasing index, p = 0.5; mean latency until initiation, p = 0.6).

**Figure 5.**
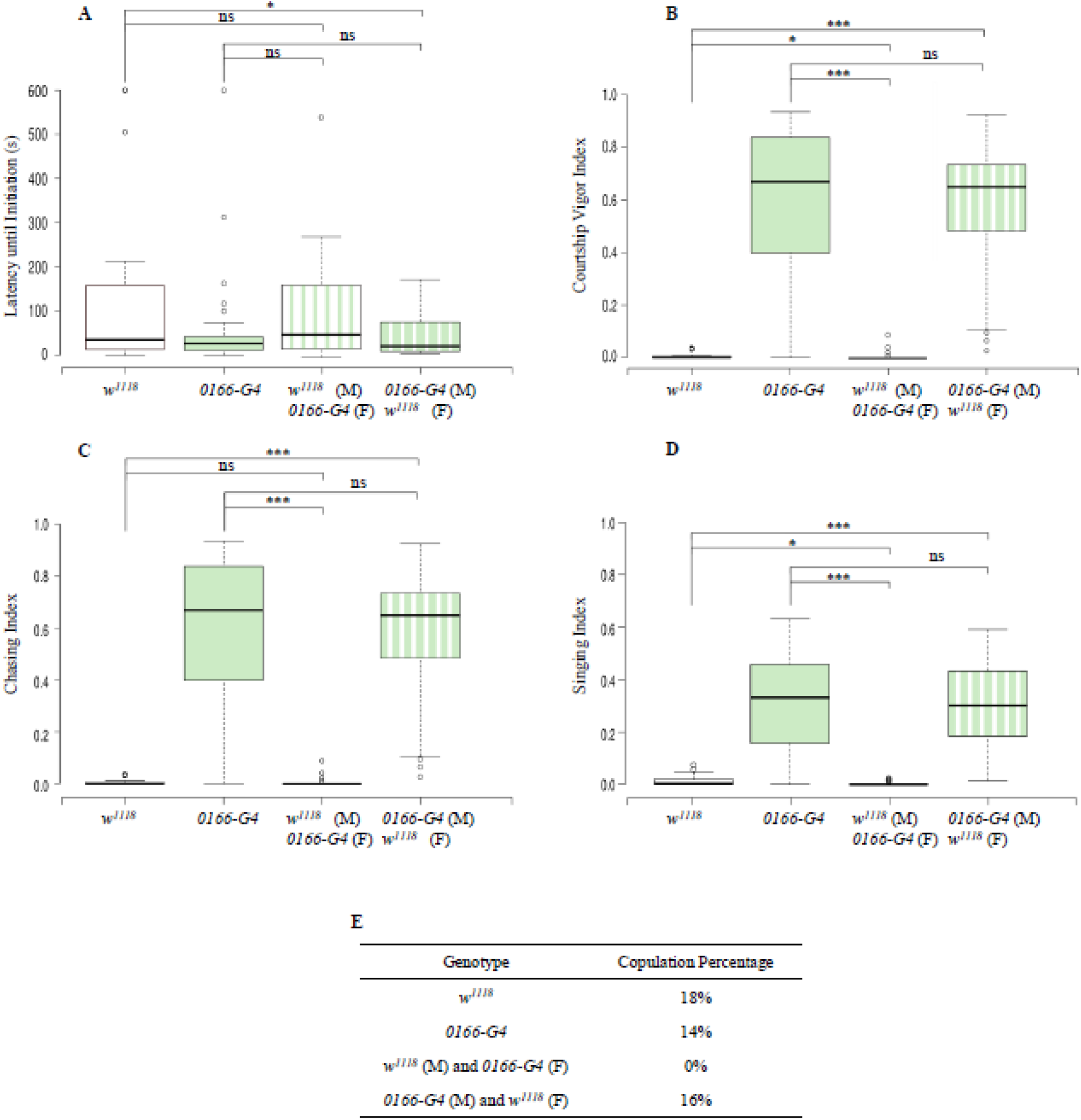
Courting intensity is affected by the male’s *Fem-1* allele and copulation percentage is affected by the female’s *Fem-1* allele copulation. (A) There is little change in the mean latency to initiation between any of the groups. (B) There is a significant increase in mean courtship vigor index between *w^1118^* and *w^1118^* males / *0166-G4* females, but no change between *0166-G4* and *0166-G4* males / *w^1118^* females. (C) Cross genotype groups show no change in mean chasing index from their respective male genotype pairs. (D) There is a significant decrease in mean singing index between *w^1118^* and *w^1118^* males / *0166-G4* females, but no change between *0166-G4* and *0166-G4* males / *w^1118^* females. (E) There is a significant reduction in copulation percentage between *w^1118^*and *w^1118^* males / *0166-G4* females, but no change between *0166-G4* and *0166-G4* males / *w^1118^* females Intensities and indices were calculated as in Figure 3, see the Methods for details. For *w^1118^*, n = 28; *0166-G4*, n = 29; *w^1118^* male / *0166-G4* female, n = 17; *0166-G4* male / *w^1118^* female, n = 19 pairs.

While the *Fem-1* females did not affect the courtship of male flies, the mutant female may have affected copulation success. The percentage of *w^1118^* female / *0166-G4* male pairs copulated was largely unchanged, yet the *w^1118^* male / *0166-G4* female pairs never copulated during the 10min observations (Figure 5E). Both the *w^1118^* and *0166-G4* control pairs copulated more frequently than the *w^1118^* male / *0166-G4* female pairs (compared to *w^1118^*, p = 0.01; compared to *0166-G4*, p = 0.01), suggesting that female receptivity may be reduced from control levels. The MATLAB analysis of path length revealed that *0166-G4* male / *w^1118^* female courting pairs did not move more or less than *0166-G4* control pairs, but that *w^1118^* male / *0166-G4* female pairs moved significantly more than *w^1118^* controls (male, p = 0.02; female, p = 5×10^−10^). This latter finding might indicate that *Fem-1* mutant females have decreased receptivity due to an overall increased level of movement that is not matched by the control males, leading to normal levels of copulation in mutant pairs and no copulation when the mutant female is courted by a control male.

### Fem-1 alleles show alterations in body and wing size of adult flies

As discussed above, *Fem-1* has been implicated in the insulin signaling pathway of mammals (Lu et al. 2005). In *Drosophila*, alterations in environmental factors or genetic manipulations of the insulin signaling pathway can result in changes in body and wing size (Oldham et al. 2002; Mirth & Shingleton 2012). We therefore used a MATLAB script to measure body length and wing length/area in the two *Fem-1* mutant flies in comparison to control flies. Body areas were not measured due to difficulties in consistently tracing the body outline in photos with variable lighting conditions. We first standardized the rearing conditions and fly age (see Methods), as growth is known to decrease in over-crowded conditions (Pitnick & García-González, 2002). The two *Fem-1* alleles showed different effects on body and wing size (Figure 6). Male *EP 2065* flies had smaller body lengths compared to w^1118^ males (p=3×10^−6^), but there were no differences in body length between female *EP 2065* and *w^1118^* flies. The *0166-G4* allele showed a large increase in female body length (p=5×10^−16^), but no change in males compared to *w^1118^*. Both sexes of *EP 2065* had smaller wing lengths than *w^1118^* flies (male, p=4×10^−5^; female, p=3×10^−7^), while there were no significant differences between the wing lengths of *0166-G4* and *w^1118^* flies. The *EP 2065* and *0166-G4* alleles both had smaller wing areas compared to *w^1118^* flies (*EP 2065:* male, p=3×10^−11^; female, p=2×10^−4^; *0166-G4*: male, p=1×10^−6^; female, p=0.05).

**Figure 6.**
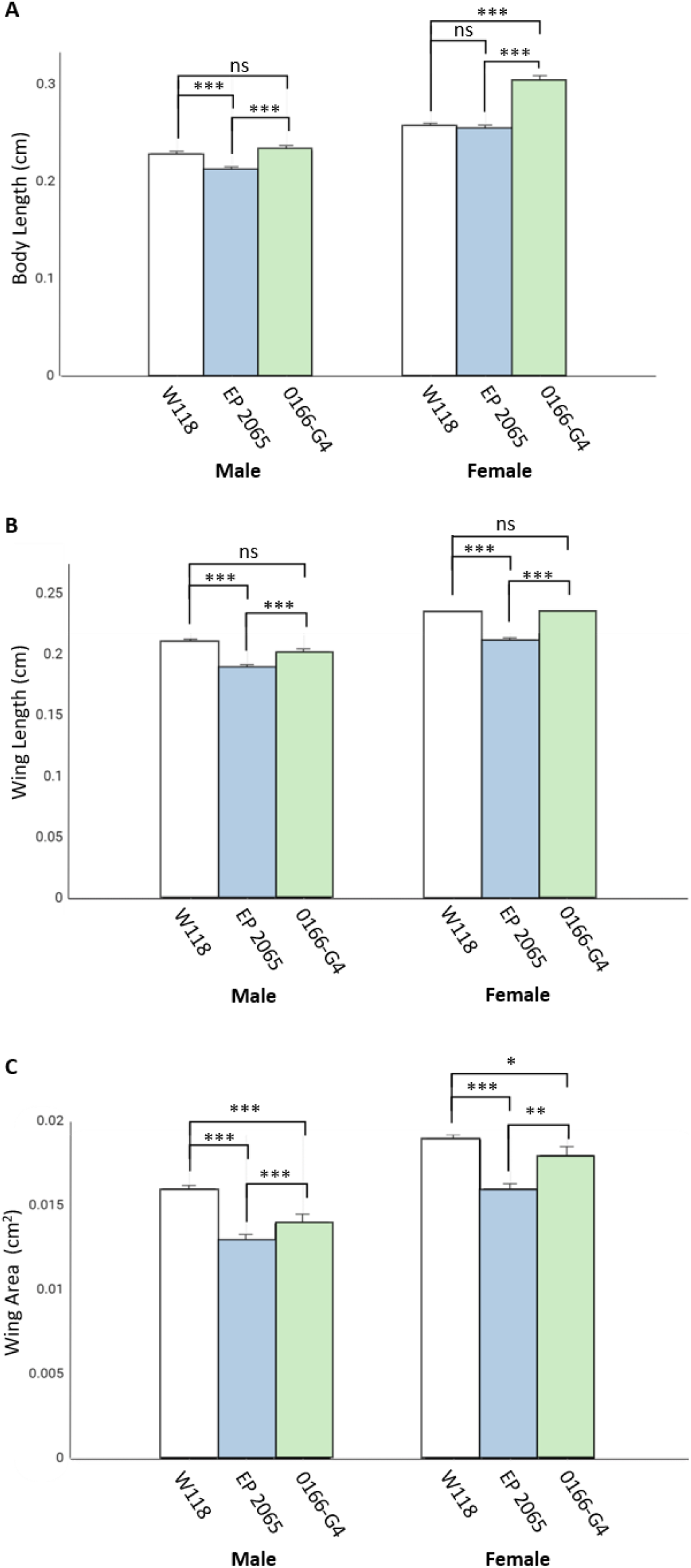
*Fem-1* mutations alter the body and wing size of adult flies. (A) Female *0166-G4* flies had significantly longer bodies than female *w^1118^* flies, while there were no differences between male *0166-G4* and *w^1118^* flies. There were no differences in body length between female *EP 2065* and *w^1118^* flies, but male *EP 2065* flies had shorter bodies than male *w^1118^* flies. For *w^1118^*: female, n = 19; male, n = 20; *EP 2065:* female, n = 20; male, n = 20; *0166-G4*: female, n = 19; male, n = 18. (B) *EP 2065* flies have significantly shorter wings than *w^1118^* flies, while there is no difference in wing length between *0166-G4* and *w^1118^* flies. For *w^1118^*: female, n = 18; male, n = 15; *EP 2065:* female, n = 20; male, n = 20; *0166-G4*: female, n = 17; male, n = 19. (C) Both *Fem1* alleles have smaller wing areas compared to *w^1118^*. For *w^1118^*: female, n = 19; male, n = 20; *EP 2065:* female, n = 20; male, n = 20; *0166-G4*: female, n = 19; male, n = 18.

## Discussion/Conclusion

This is the first study to investigate a role for the *Fem-1* gene in *Drosophila*. We analyzed two independent *Fem-1* alleles and demonstrate that these alleles overlap in their courtship phenotypes but differ in the extent of these phenotypes (Figures 3 and 5). Considering the importance of *Fem-1* in the sexual development of *C. elegans* and other mammals, it was expected that *Fem-1* might influence courtship behavior in *Drosophila*. Cross-genotype experiments demonstrated that the *Fem-1* mutations in the male predominantly determine courting intensity (Figure 3), while the female *Fem-1* allele may affect copulation success (Figure 5). Similarly, the size differences we observed in the adult wing and body might suggest a role for *Fem-1* in tissue growth (Figure 6), consistent with potential effects on insulin signaling as is observed in mice (Lu et al., 2005; Mirth & Shingleton, 2012; Oldham et al., 2002). To fully characterize the role of *Fem-1* in *Drosophila* courtship and tissue development, more alleles should be investigated using specialized genetic tools (that don’t yet exist) such as targeted knockdown of the gene in small subsets of cells and tissues.

Our analysis of *Fem-1* mutants showed that it functions in an evolutionarily conserved manner. We chose these assays based on the assumption that *Drosophila* Fem-1 would function similar to Fem-1 in other organisms. The experiments used the available fly strains, and it should be noted that the two alleles have not been assayed for effects on mRNA expression or protein function. We also did not clean up their genetic backgrounds. Nevertheless, both alleles showed largely similar phenotypes, especially with regard to courtship, which forms the basis of our discussion. Beyond courtship and growth, *Drosophila* provides a wealth of other experimental paradigms with which to examine Fem-1 function, including several adult and larval behaviors and their underlying neural circuits and a detailed analysis of synapse development at the larval neuromuscular junction (Broadie & Bate, 1995).

### Fem-1 may affect sexual development in Drosophila

Although the upstream regulatory proteins in the *Drosophila* sex determination cascade are well studied (Pomiankowski, Nöthiger, & Wilkins, 2004), there remain many unidentified downstream effectors such as regulators of *Sex-lethal*, regulation by circadian rhythms, and actions of neuromodulators (Ellendersen & von Philipsborn, 2017; Kim et al., 2017, Fujii & Amrein 2002; Salz & Erikson 2010; Fujii et al., 2017). Considering the striking change in courtship behavior in *Fem-1* mutants, it is possible that *Fem-1* interacts with components of the sex determination pathway. *Fem-1* has a well-defined role in *C. elegans* sex determination, where *Fem-1* helps to degrade *transformer-1* in male worms (Starostina et al. 2007). However, large differences exist between the genetic mechanisms of sex determination of flies and worms. Sex determination in *Drosophila* is mediated by a cascade of regulated mRNA splicing (Haag & Doty 2005). *Sex-lethal (Sxl)* is the genetic switch that determines male or female development in *Drosophila* by regulating the mRNA splicing of the female-specific, *Drosophila* homolog of *transformer-1* (Salz & Erickson, 2010). Given that both *Fem-1* proteins (Fem-1a and Fem-1b) contain ankyrin-repeats, it is very likely that their interactomes could be identified by pulldown or gel-shift assays once an antibody to Fem-1 is created. An antibody against mouse Fem-1b already exists (Lu et al. 2005) and could be useful if it cross reacted with one or both of the transcripts in *Drosophila*.

Drawing comparisons between *Fem-1* mutant phenotypes in *Drosophila* and other sex determination mutants could also be useful. Male courtship behaviors are dependent upon the splicing of the *fruitless* (*fru*) gene, which is differentially spliced between males and females (Demir & Dickson 2005). Mutations that decrease the expression of male-specific *fru* result in decreased courting intensity and null mutations completely disrupt male courting (Anand et al. 2001). *Fem-1* mutants have male-specific phenotypes, so future investigations of *fru* expression and *fru / Fem-1* double mutants may indicate protein-protein interactions that affect *fru* signaling. In addition, the *found-in-neurons* (*fne*) gene encodes an RNA binding protein whose loss of function shows decreased courting intensity, mating frequency, and axonal pathfinding errors during the development of the CNS (Zanini et al. 2012). Double mutant experiments on *Fem-1 / fne* could therefore give insight into *Fem-1’s* effect on courting intensity and potential roles in neuronal development. If an RNAi transgene were to be made for *Fem-1*, then tissue-specific expression tools could be used to knock down *Fem-1* in elements of the courtship circuitry (von Philipsborn et al. 2011). This would give more insight into what part of the courtship circuit is affected by *Fem-1*.

### Fem-1 may contribute to tissue growth in Drosophila

*Fem-1* studies in mice have given valuable insight into its role in insulin signaling. *Fem-1* mutations alter the secretion of insulin by pancreatic cells (Lu et al. 2005). The sexual development and insulin secretion defects seen in *Fem-1* mice may be linked in some way, considering that insulin receptors are crucial for genital development and primary sex determination in the mouse (Pitetti et al. 2013). In *Drosophila*, sex determination and insulin signaling could also be linked, since insulin mediates sexual attractiveness (Kuo et al. 2012). While we did not measure insulin levels or insulin receptivity in the *Fem-1* alleles, we did document subtle changes in fly and wing size (Figure 6). It may therefore be useful to examine these *Fem-1* phenotypes in more detail and perhaps in combination with mutations that affect insulin production or signaling.

We have presented a preliminary analysis on the effects of *Fem-1* mutation on courtship, primary sexual characteristics, and growth in *Drosophila*. To characterize the precise role of *Fem-1* in sexual determination and insulin signaling, further study should be done using specialized genetic tools.

## Abbreviations

EP 2065: **enhancer-promoter insert 2065**
*0166-G4*: **Fem-1^0166-G4^**
*w*^*1118*^: *white mutation, allele 1118*, **genetic control**

## Acknowledgements

We would like to thank the Berke Laboratory for their constructive comments and the Biology Department at Truman State University for funding this project through Dr. Berke’s start-up funds and a TruScholar award to M. Thies.

